# Distinct Neurogenic Dynamics of Cortico-cortical Neuronal Subtypes in Layer 2/3 of the Mouse Visual Cortex

**DOI:** 10.1101/2025.07.17.665445

**Authors:** Mustapha Major, Paul Pham, Efrain Hernandez-Alvarez, Euiseok J. Kim

**Affiliations:** Department of Molecular, Cell, and Developmental Biology, University of California, Santa Cruz, CA 95064, USA; Institute for the Biology of Stem Cells, University of California, Santa Cruz, CA 95064, USA

**Keywords:** neurogenesis, cortico-cortical projection neurons, L2/3 neurons, mouse visual cortex, birthdating, neuronal subtypes

## Abstract

In the mammalian cerebral cortex, the birthdates of excitatory projection neurons are closely linked to their laminar positions, which are often associated with distinct long-range projection targets. Although broad relationships between neurogenic timing, laminar position, and projection patterns are well established, the degree to which birthdate specifies projection identity within the same cortical layer remains unclear. In the mouse primary visual cortex (V1), neurons projecting to lateral higher visual areas are relatively evenly distributed throughout layer 2/3, whereas those projecting to medial areas are biased toward more superficial sublayers. To determine whether these projection identities are linked to neurogenic timing, we combined EdU birthdating with retrograde viral tracing. Notably, we found that V1 layer 2/3 neurons projecting to lateral higher visual areas are preferentially born at embryonic day 15.5 (E15.5) compared to E16.5, whereas V1 neurons projecting to medial higher visual areas show no significant birthdate bias between E15.5 and E16.5. These findings suggest that distinct cortico-cortical projection subtypes in layer 2/3 are generated on different temporal schedules, linking neurogenic timing to fine-scale projection identity.

## INTRODUCTION

Understanding how diverse neuronal subtypes emerge during development is a central question in neurobiology. The mammalian brain consists of a wide array of cell types distinguished by morphology, molecular identity, physiology, connectivity, and anatomical location. Historically, neurons were classified by gross morphology and anatomical location (Glickstein, 2006; Nelson et al., 2006; Fishell and Heintz, 2013), but recent advances in transcriptomics and circuit mapping have enabled much finer classifications, revealing hierarchical taxonomies of neuronal types across the mouse brain (Tasic et al., 2016; Zeng and Sanes, 2017; Yao et al., 2021). Within these taxonomies, excitatory neurons in the neocortex are broadly classified into major projection classes such as intratelencephalic (IT or cortico-cortical projection neurons, CCPNs), extratelencephalic (ET), corticothalamic (CT), and near-projecting types based on their laminar position and long-range connectivity (Harris and Shepherd, 2015).

However, even within a single layer and projection class, further subclassification reveals considerable cellular diversity. For example, in layer 2/3 (L2/3) of the mouse primary visual cortex (V1), CCPNs project to a variety of higher visual areas (HVAs) with specific projection motifs forming anatomical subtypes (Wang and Burkhalter, 2007; Han et al., 2018; Kim et al., 2020). Among these, neurons projecting to the anterolateral (AL) and posteromedial (PM) HVAs, L2/3 V1→AL and L2/3 V1→PM respectively, form largely non-overlapping groups with distinct properties. They differ in visual tuning preferences (Glickfeld et al., 2013) and local circuit connectivity (Kim et al., 2018), rarely co-project to both AL and PM (Kim et al., 2018, 2020), and receive selective long-range feedback predominantly from their respective target areas (Kim et al., 2020). These features support the notion that L2/3 CCPNs in V1 comprise discrete subtypes with distinct circuit identities.

One underexplored question is how these subtype-specific projection identities are developmentally specified. A clue may lie in their laminar positioning: L2/3 V1→AL neurons are evenly distributed across L2/3, whereas L2/3 V1→PM neurons are more superficial (Kim et al., 2018, 2020; Cheng et al., 2022; Han and Bonin, 2024). Since cortical neurons are generated in an inside-out sequence, with earlier-born neurons settling deeper and later-born neurons migrating to more superficial positions (Angevine and Sidman, 1961; McConnell, 1995; Rakic, 2007), we hypothesized that birthdate contributes to projection identity within L2/3.

To test this hypothesis, we asked whether V1 neurons projecting to lateral versus medial HVAs differ in their birth timing. We defined AL, rostrolateral (RL), and lateromedial (LM) as lateral HVAs (L-HVAs) and PM and anteromedial (AM) as medial HVAs (M-HVAs), based on their axonal projection targets (Figure 1A). To assess birthdate, we administered EdU at embryonic days E14.5, E15.5, and E16.5, and used fluorescently tagged AAVretro virus tracers to label V1 CCPNs projecting to either L-HVAs or M-HVAs. By comparing the birthdate distributions of these projection-defined populations, we determined whether distinct L2/3 CCPN subtypes arise at different developmental time points.

**Figure 1.**
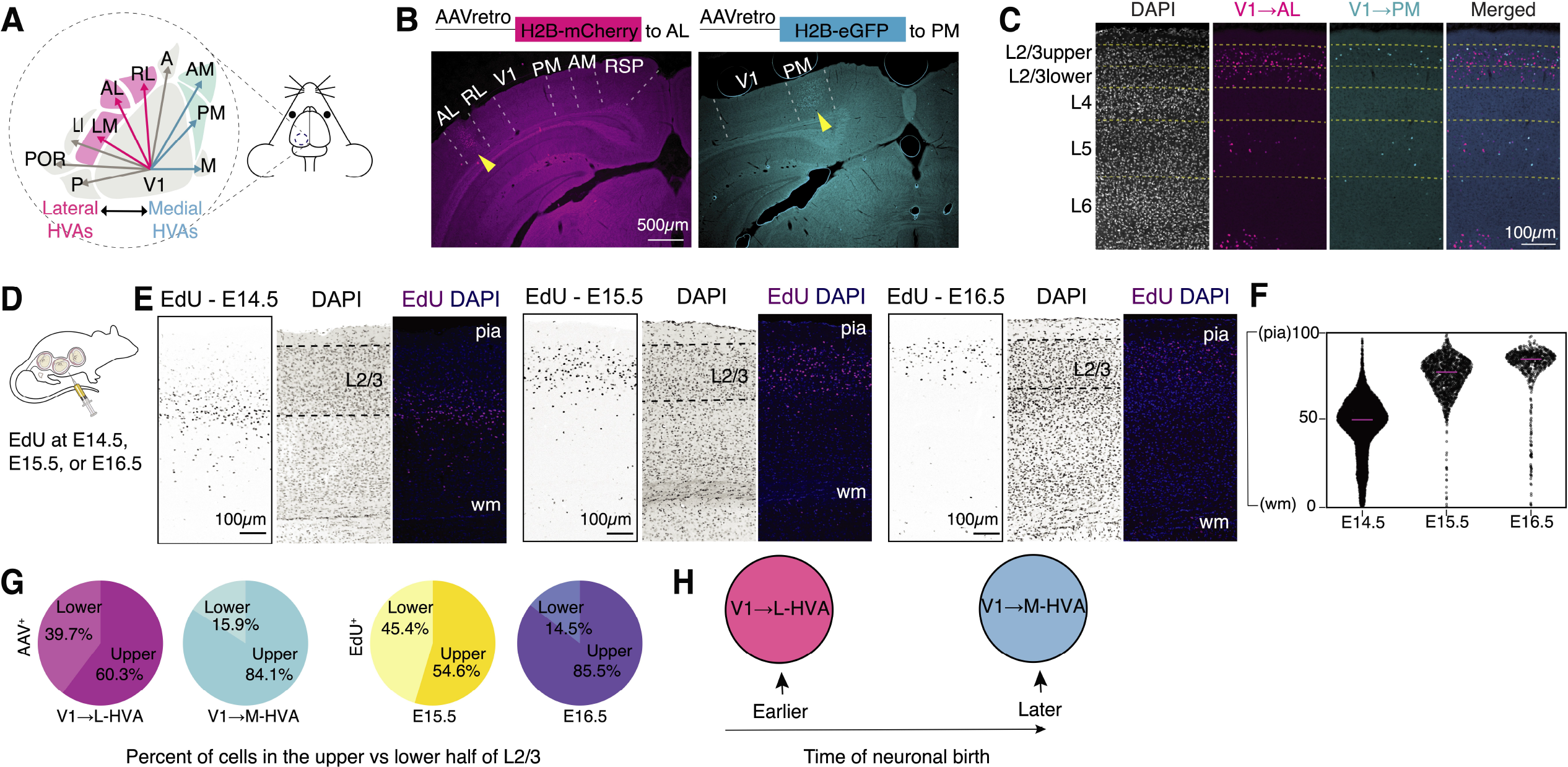
Differential laminar distribution of V1 neurons projecting to distinct higher visual areas (HVAs) and of EdU-labeled neurons from E14.5 to E16.5. **(A)** Schematic of primary visual cortex (V1) and surrounding HVAs. **(B)** Coronal sections showing representative retrograde labeling using AAVs injected into lateral (AL, left) and medial (PM, right) HVAs, with yellow arrowheads indicating the injection sites. **(C)** Example coronal images of AAVretro-labeled neurons in V1 following injections into AL (magenta) and PM (cyan). **(D)** Diagram illustrating the EdU labeling protocol during embryonic development. **(E)** Representative images of EdU-labeled cells in coronal V1 sections from mice injected at E14.5 (left), E15.5 (middle), or E16.5 (right) and analyzed in adulthood. **(F)** Violin plots showing the distribution of EdU+ cells along the cortical depth from pia to the white matter (wm) in three example adult mice across the whole primary visual cortex. Data are represented as median. **(G)** Pie charts illustrating the proportion of AAV+ (left) and EdU+ (right) neurons located in the upper versus lower half of V1 L2/3. Data are represented as mean. **(H)** Working model: L2/3 V1 neurons projecting to lateral HVAs (L-HVAs) are predominantly born earlier, whereas those projecting to medial HVAs (M-HVAs) are born later.

## METHODS

### Experimental animals and husbandry

C57BL/6J mice were used as wild-type. Both male and female mice were used. The specific ages and conditions of the experimental animals are described in Table 1. All mice were housed with a 12-hour light and 12-hour dark cycle and *ad libitum* access to food and water. All animal procedures were performed in accordance with the University of California, Santa Cruz animal care and use committee (IACUC)’s regulations.

**Table 1.**
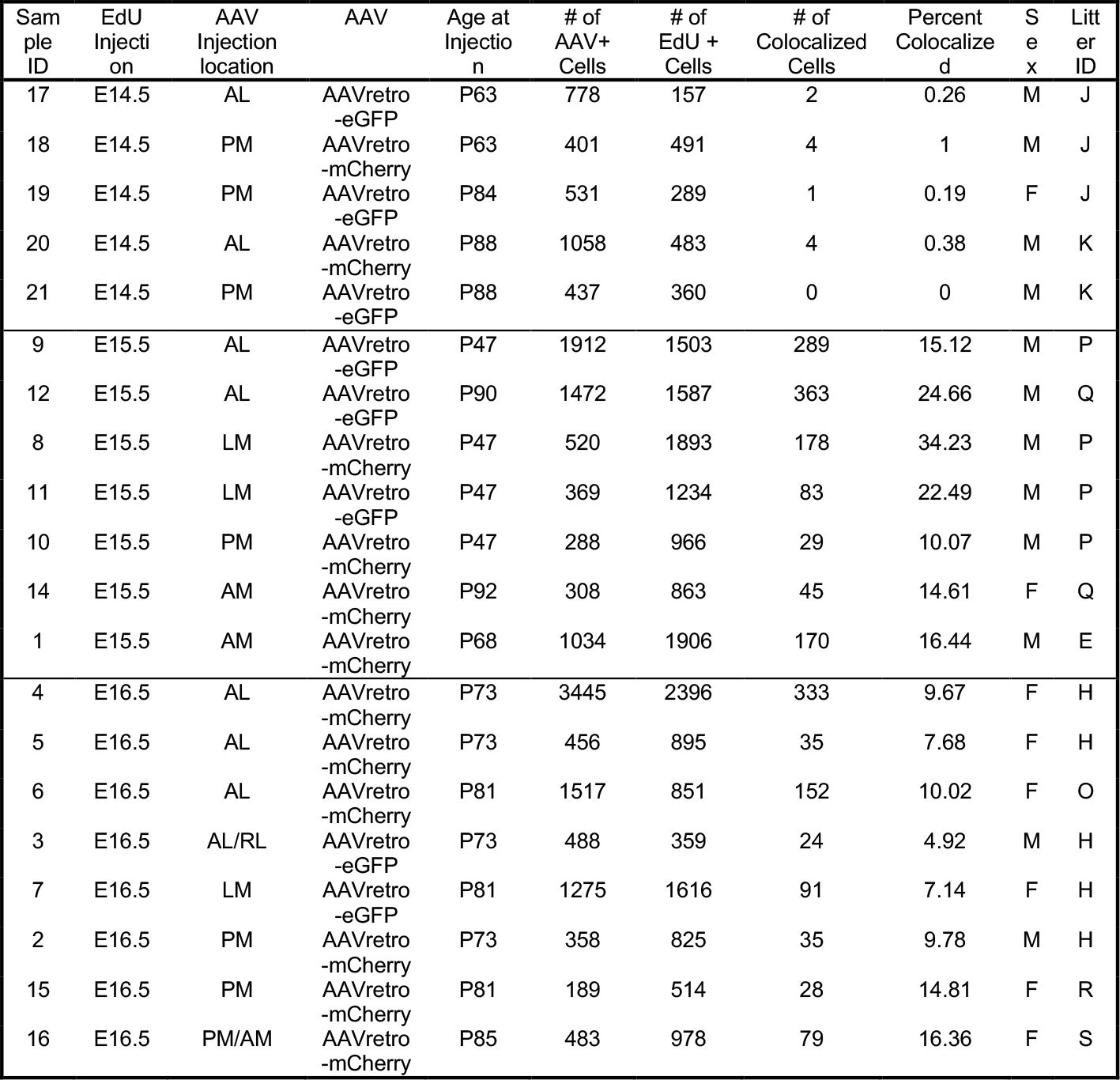
Summary of animal information, experimental conditions, cell counts and colocalization percentages.

### EdU (5-ethynyl-2’-deoxyuridine) injection

Male and female wildtype mice were crossed to each other with a 12-hour breeding window, and females were checked daily for vaginal plugs to determine date of conception. The time point at which a plug was detected was considered embryonic day (E) 0.5. Pregnant dams were given one intraperitoneal (IP) injection of 25mg/kg EdU (MilliporeSigma, 900584) at E14.5, E15.5, or E16.5.

### Virus preparation and stereotaxic animal surgery

Adeno-associated virus (AAVs) were produced by the Salk Viral Core GT3: scAAVretro-hSyn-H2B-eGFP (referred to hereafter as AAVretro-eGFP, 1.16×10^13^ GC/ml), scAAVretro-hSyn-H2B-mCherry (referred to hereafter as AAVretro-mCherry, 1.21×10^13^ GC/ml). Mice from postnatal day (P) 47 to P92 were used. Mice were anesthetized with an IP injection of a cocktail containing 80.4 mg/kg ketamine and 8 mg/kg xylazine cocktail, and continuous inhalation of 1-3% gaseous isoflurane throughout the procedure. To label V1 neurons projecting to AL, PM, and other HVAs, 15-50 nl of AAVretro-eGFP or AAVretro-mCherry was injected into the target HVA. The fluorescent protein used in the lateral or medial injection was alternated between each animal. Injection sites were located using stereotaxic coordinates relative to lambda on the mediolateral and anteroposterior axes, and relative to the pia on the dorsoventral axis: 3.5 mm lateral, 1-1.5 mm anterior, and 0.3 mm ventral for AL or L-HVAs; 1.6 mm lateral, 1.23-1.6 mm anterior, and 0.3-0.4 mm ventral for PM or M-HVAs. A drilled burr hole was made over the target area, and injections were done with a 20-30 μm diameter glass pipette, using air pressure from a 1 ml syringe with 18G tubing adapter and tubing. Post-surgery, 5 mg/kg of carprofen was injected intramuscularly, and mice were given water with ibuprofen (30 mg/kg).

### Histology

Ten days after virus injection, mice underwent trans-cardiac perfusion using 1X phosphate-buffered saline (PBS) containing heparin (10 U/ml; Sigma-Aldrich, H3393) followed by 4% paraformaldehyde (PFA). Brains were dissected from skulls and postfixed with 2% PFA and 15% sucrose in PBS at 4°C overnight and then immersed in 30% sucrose in 1X PBS at 4°C for a minimum of 24 hours before sectioning. Using a sliding microtome (Espredia, HM430), 50 μm coronal brain sections were cut and stored in 1X PBS with 0.01% sodium azide or cryogen (3 parts ethylene glycol, 3 parts glycerol, 3 parts ddH_2_O, 1 part 10x PBS) at 4°C. After washing 3 times, 10 minutes each with 1X PBS, EdU staining solution (100 mM Tris pH8.0, 4 mM CuSO_4_, 0.65 μM sulfo-Cy5 azide, 10 mM sodium ascorbate) was applied and incubated for 30 minutes at room temperature, protected from light. Sections were washed three times for 10 minutes each in 1X PBS, then incubated with DAPI (4′,6-diamidino-2-phenylindole; ThermoFisher Scientific Invitrogen, D1306) for 30 minutes at room temperature, protected from light. After an additional three washes in 1X PBS (10 minutes each), sections were mounted on slides using a polyvinyl alcohol mounting medium containing DABCO (PVA-DABCO) and allowed to air dry.

### Imaging and quantification

Brain sections were imaged using a 4×/0.2 NA (wd: 20 mm) or a 10×/0.45 NA (wd: 4 mm) objective on a Keyence BZ-9000 microscope. For each animal, injection sites were evaluated to confirm that labeling was restricted to the appropriate cortical visual areas. Representative images were acquired using a Zeiss LSM 880 confocal microscope with either a 10×/0.45 NA (wd: 2.0 mm) or a 40×/0.95 NA corr (wd: 0.25 mm) objective. To quantify single- and double-labeled neurons in V1, cortical borders were delineated on each section based on the Allen Mouse Brain Common Coordinate Framework Reference Atlas (Wang et al., 2020), and L2/3 was manually defined using the DAPI nuclear counterstain based on cell density. AAVretro-labeled neurons (AAV+) and double-labeled AAV+EdU+ neurons were manually counted. Because EdU signal intensity diminishes with each successive cell division (Pereira et al., 2017), only strongly labeled cells defined as having a mean intensity greater than 50% of the brightest EdU+ cell within a given section were included in the analysis to select for neurons born near the time of EdU injection. Single-labeled EdU+ neurons were automatically detected using QuPath (Bankhead et al., 2017) and custom scripts applying the same intensity threshold.

### Statistical analysis

P-values were calculated using either Student’s t-test, the Kruskal–Wallis test, or the Pearson correlation test with Python 3.12, depending on the distribution and characteristics of the data. Normality and equality of variance were assessed prior to statistical testing using the Shapiro-Wilk and Levene’s test, respectively.

## RESULTS

To examine the sublaminar distribution of L2/3 CCPNs targeting medial (e.g., AM or PM) versus lateral (e.g., AL, LM, or RL) higher visual areas (HVAs) in the adult mouse cortex (P47– P92), we injected AAVretro-eGFP and AAVretro-mCherry into medial and lateral HVAs, respectively. Brain tissue was collected 10 days post-injection for analysis (Figures 1A-B). Coronal brain sections containing the visual cortex were assessed using the Allen mouse brain common coordinate framework to confirm accurate targeting of the retrograde AAVs to higher visual areas (Figure 1B) (Wang et al., 2020). Fluorescence imaging revealed that V1 CCPNs projecting to lateral and medial visual areas were differentially distributed within L2/3. Specifically, PM- or AM-projecting V1 CCPNs (V1→M-HVAs) were concentrated at 28.19 ± 2.61% of the normalized depth within layer 2/3, measured from the top of the layer. In contrast, AL-, RL- or LM-projecting V1 CCPNs (V1→L-HVAs) were more evenly distributed through the depth of L2/3 with the average depth 43.59 ± 1.34% (Figures 1C). This distribution pattern aligns with previous reports (Kim et al., 2018, 2020; Han and Bonin, 2024), including those utilizing alternative retrograde tracers such as cholera toxin subunit B (Kim et al., 2020).

L2/3 neurons in the mouse cortex are predominantly generated between embryonic days (E) 14.5 and 16.5 (Vitali et al., 2018; Baumann et al., 2025). To assess neuronal birthdates within the visual cortex, we administered EdU, a thymidine analog, to pregnant dams at E14.5, E15.5, or E16.5 (Figure 1D). Although some EdU+ cells labeled at E14.5 were detected in L2/3, the majority localized to layer 4 (Figures 1E-F). In contrast, EdU+ cells labeled at E15.5 and E16.5 were abundantly present in L2/3 (Figures 1E-G). Among these, cells labeled at E15.5 were positioned significantly deeper from the pial surface than those labeled at E16.5 (48.10 ± 2.49%, 27.38 ± 1.73%, Figure 2).

**Figure 2.**
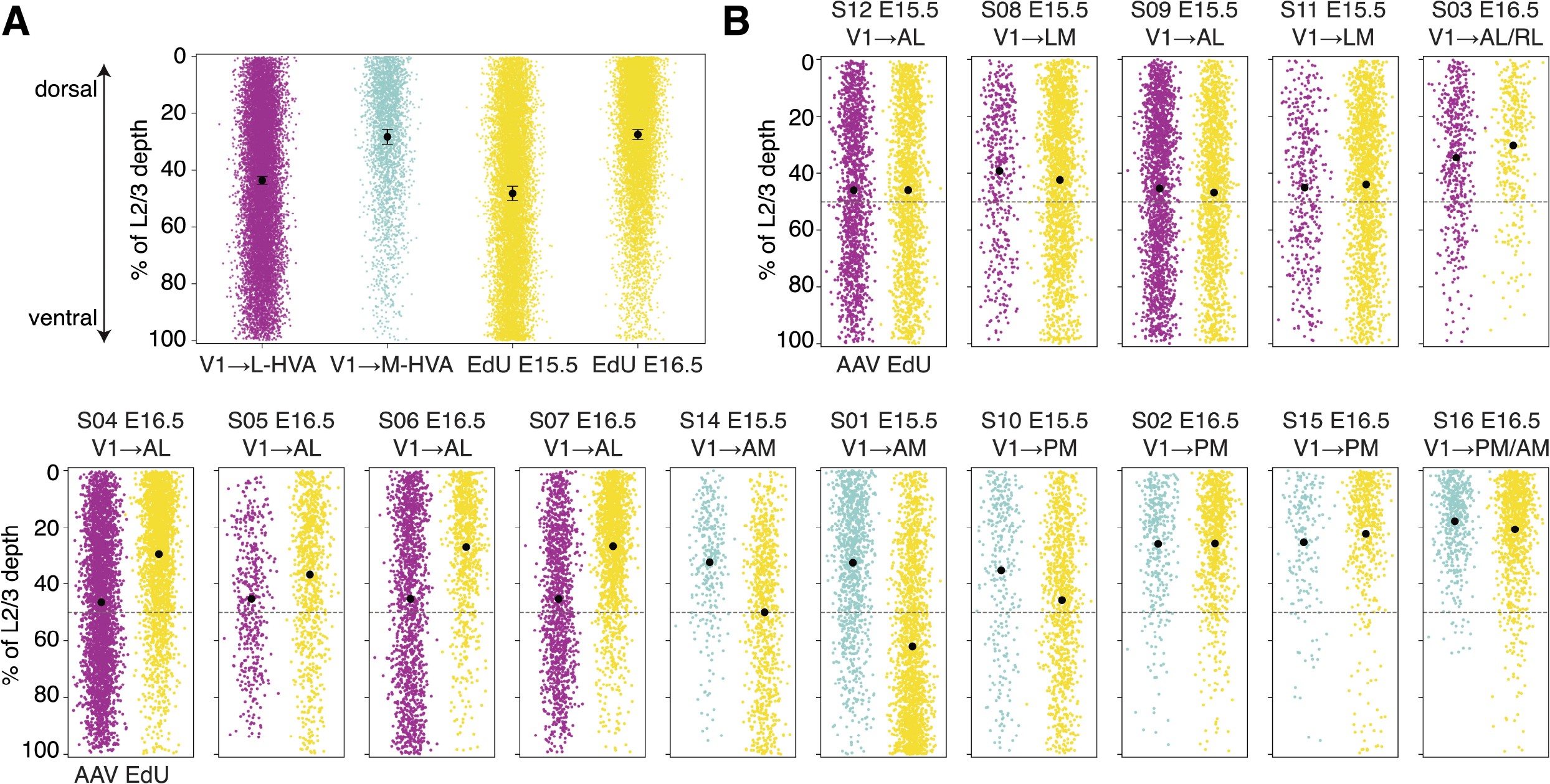
Sublaminar positioning of L2/3 V1→L-HVA and V1→M-HVA neurons parallels the laminar distributions of neurons born at E15.5 and E16.5. **(A)** Scatter plots showing the distribution of soma depth for AAV-labeled and EdU-labeled cells, measured relative to the dorsal boundary of layer 2 (0%) and the ventral boundary of layer 3 (100%). Black points indicate mean and error bars indicate SEM. **(B)** Scatter plots showing soma depth distributions of individual AAV+ and EdU+ cells from individual animals, using the same reference points as in **(A)**. The dashed line indicates the midpoint of L2/3 depth.

We further quantified the proportion of neurons located in the upper versus lower half of L2/3. 39.68% of V1→L-HVA CCPNs resided in the lower half, compared to only 15.90% of V1→M-HVA CCPNs. Likewise, 45.35% of E15.5-born neurons were located in the lower half of the layer, while only 14.50% of E16.5-born neurons showed similar positioning (Figures 1G and 2). Both the mean and median soma depths were deeper for V1→L-HVA neurons (mean: 43.59%, median: 41.77%) than for V1→M-HVA neurons (mean: 28.19%, median: 23.59%), a pattern also reflected in the E15.5 (mean: 48.10%, median: 46.49%) versus E16.5 (mean: 27.38%, median: 22.66%) groups (Figure 2). This pattern was consistent among all individual samples with the exception of sample ID 01, which exhibited an EdU+ distribution that was intermediate to stereotyped E14.5 and E15.5 distribution patterns (Figure 2B).

Given the resemblance between the soma distribution of AL- or laterally projecting V1 CCPNs and that of E15.5-born L2/3 neurons, and between PM- or medially projecting V1 CCPNs and E16.5-born L2/3 neurons, we hypothesized that laterally projecting V1 CCPNs are predominantly generated at E15.5, whereas medially projecting CCPNs originate mostly at E16.5 (Figure 1H). To test this, we quantified the proportion of EdU+ neurons among AAVretro-labeled laterally or medially projecting V1 CCPNs within L2/3 across the entire V1 (Figure 3). As expected, EdU incorporation at E14.5 resulted in nearly zero colocalization with AAVretro-labeled CCPNs, regardless of projection target, reflecting the low number of E14.5-born neurons in L2/3 (Figure 3B). For labeled V1→L-HVA CCPNs, the percentage of EdU+ neurons was significantly higher in the E15.5 group than in the E16.5 group (24.12 ± 3.94% and 7.89 ± 0.92%, respectively), supporting our hypothesis. This trend remained consistent in animals with AAVretro injections specifically targeted to AL. In contrast, for V1→M-HVA CCPNs, no significant difference in colocalization was observed between the E15.5 and E16.5 groups (13.71 ± 1.89% and 13.65 ± 1.99% respectively, Figure 3B).

**Figure 3.**
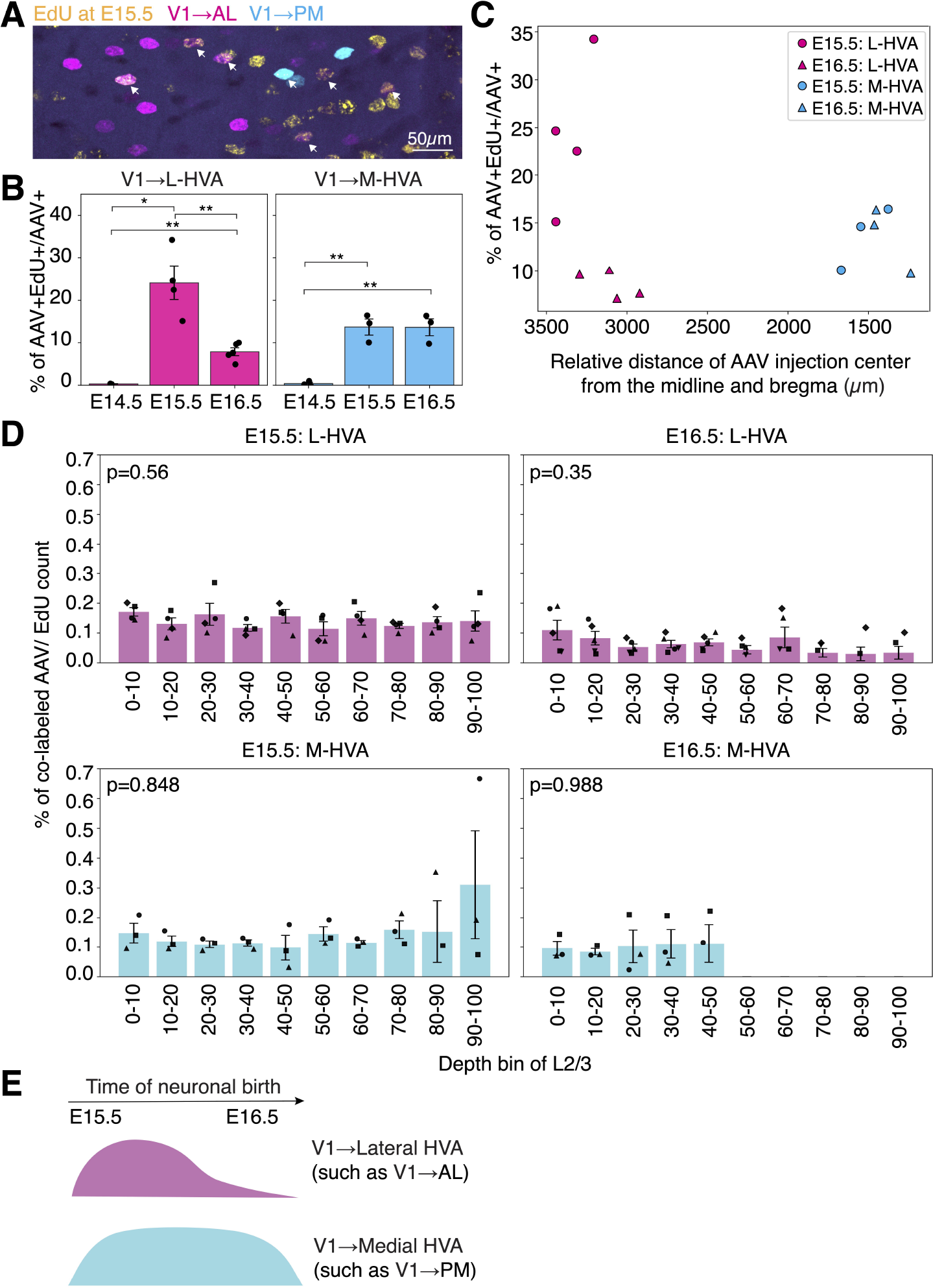
L2/3 V1→L-HVA neurons are born in a greater proportion at E15.5 than at E16.5, while L2/3 V1→M-HVA neurons show no temporal bias. **(A)** Confocal image of L2/3 in a coronal section of V1. AAV-labeled neurons projecting to AL (magenta) and PM (cyan) are shown alongside EdU+ neurons labeled at E15.5 (yellow). Arrows indicate cells with colocalized AAV and EdU signals. **(B)** Percentage of AAV+ cells that are also EdU+ at E14.5, E15.5, or E16.5 in animals injected with AAVretro in a lateral HVA (left) or medial HVA (right). Each dot represents an individual animal. Student’s t-test; *p<0.05, **p < 0.01. **(C)** Colocalization percentage plotted against the distance of the center of the AAVretro injection site from the midline and anterior position of the brain. **(D)** Normalized laminar distribution of AAV+EdU+ neurons across L2/3 for each injection group. Kruskal-Wallis test. **(E)** Model summarizing temporally distinct neurogenesis of V1 L2/3 cortico-cortical projection neurons targeting lateral versus medial HVAs. n=2 for E14.5 L-HVA, n=3 for E14.5 M-HVA, n= 4 for E15.5 L-HVA, n= 3 for E15.5 M-HVA, n=5 for E16.5 L-HVA, and n=3 for E16.5 M-HVA. Data are represented as mean ± SEM.

To determine whether AAV and EdU colocalization rates were influenced by the spatial location of AAVretro injection sites, we analyzed the relationship between colocalization percentages and the medial–lateral and anterior–posterior coordinates of injection centers relative to bregma (Figure 3C). For lateral injections, there was a slight negative trend between colocalization percentage and distance from both the midline and anterior visual cortex; however, these correlations were not statistically significant (p = 0.225, Pearson correlation test, Figure 3C). Similarly, no significant correlation was observed between colocalization percentage and spatial coordinates for medial injections (Figure 3C). To determine whether the percentage of AAV and EdU colocalization was influenced by EdU+ cell density or laminar position within L2/3, we divided the full depth of L2/3 into ten equally spaced bins and quantified the proportion of AAV+EdU+ neurons in each bin, normalized to the local EdU+ cell density (Figure 3D). This analysis was performed across all four experimental groups. We found no statistically significant differences in the distribution of colocalization percentages along the L2/3 depth in any of the groups. These results suggest that differences or similarities in AAV+EdU+ colocalization rates between animal groups are not driven by confounding factors such as injection site location, EdU labeling density, or cortical depth.

## DISCUSSION

In summary, using a combination of birthdating and retrograde labeling, we found that L2/3 V1→L-HVA neurons are preferentially generated at E15.5, with reduced production at E16.5, whereas L2/3 V1→M-HVA neurons are generated at similar rates across both time points. These findings indicate a temporally biased wave of neurogenesis for L-HVA-projecting neurons, in contrast to a more uniform generation of M-HVA-projecting neurons (Figure 3E).

L2/3 CCPNs display considerable diversity in gene expression, local and long-range connectivity, and functional roles (Tasic et al., 2016; Meng et al., 2017; Kim et al., 2018, 2020; Yao et al., 2021). Our data reveal temporal differences in the birth rates of V1 CCPN subtypes. However, neuronal birthdate alone does not appear to dictate projection identity, as L2/3 CCPNs generated at the same embryonic stage can project to different targets. This raises the question of how discrete yet temporally close waves of neurogenesis give rise to such heterogeneous subtypes. In V1, the diversity among L2/3 CCPNs projecting to higher visual areas (HVAs) may stem from differences in progenitor origin: for example, direct versus indirect neurogenesis from radial glia or intermediate progenitors (Ellender et al., 2019; Huilgol et al., 2023). Alternatively, this heterogeneity could reflect depth-dependent, continuous changes in transcriptional programs that govern cell-type specification during cortical development, consistent with the birthdate-dependent inside-out pattern of laminar organization (Cheng et al., 2022; Xie et al., 2025).

Our study has several limitations. First, EdU-based birthdating provides relatively coarse temporal resolution, making it difficult to precisely determine the exact timing of neuronal birthdates (Landy et al., 2021). Mice were bred using a 12-hour mating window, with the time of plug detection designated as E0.5, introducing a potential 12-hour variability in labeling across animals. Although the spatial distribution of EdU+ cells labeled at E14.5, E15.5, and E16.5 was generally consistent across litters, one animal (sample ID 01) showed an intermediate distribution between typical E14.5 and E15.5 patterns (Figure 2B and Table 1), likely reflecting this variability. Nevertheless, we mitigated this potential limitation by analyzing multiple animals from independent breeders and litters, and by applying stringent criteria to identify birthdated neurons, selecting EdU+ cells that incorporated the label during their final S-phase before exiting the cell cycle on the day of EdU administration (See Methods). Second, retrograde labeling methods reveal only projections to the injected target site and do not capture the full extent of a neuron’s axonal arborization (Han et al., 2018; Winnubst et al., 2019). Consequently, the complete projection patterns of CCPNs were not mapped in this study, and a comprehensive analysis linking birthdate to full projection identity remains to be conducted.

Building on the current findings, future studies using complementary approaches may help clarify how birth timing contributes to the development of L2/3 neurons with distinct projection identities. One promising strategy is genetic birthdating using transgenic mouse lines that allow permanent labeling of neurons born at specific embryonic stages. For instance, inducible CreER lines such as Neurog1-CreER, Neurog2-CreER, or Tbr2-CreER, crossed with a Cre-dependent Flp-expressing reporter (e.g., loxP-STOP-loxP-Flp), can be used in combination with tamoxifen administration at defined embryonic time points (Kim et al., 2011; Hirata et al., 2021; Matho et al., 2021). In adult offspring, a Flp-dependent viral tracer such as AAV-fDIO (or FLEx^frt^)-fluorescent protein can be injected into V1 to selectively label axons of neurons born at those time points. This approach would enable direct quantification of V1 CCPN axons in medial versus lateral HVAs, allowing comparisons between early- and late-born L2/3 neurons. In parallel, transcriptomic or epigenomic profiling of these developmentally tagged neurons could uncover molecular mechanisms underlying subtype specification and projection identity (Isshiki et al., 2001; Molyneaux et al., 2007; Greig et al., 2013; Cheng et al., 2022). An important question for future investigation is whether similar temporal patterns of neurogenesis are observed in CCPNs of other cortical areas. For example, L2/3 CCPNs in the secondary visual cortex projecting to other HVAs, or those in the primary somatosensory or auditory cortices projecting to secondary cortical regions (Yamashita et al., 2018; Liu et al., 2019).

## Author Contributions

MM, PP, EH performed experiments. MM and EJK analyzed data. MM and EJK wrote the manuscript. All authors contributed to the article and approved the submitted version.

## Acknowledgements

We thank Bin Chen, Bradley Colquitt, Richard Dickson, and Matthew Jacobs for reading the manuscript. We acknowledge technical support from Benjamin Abrams, UCSC Life Sciences Microscopy Center, RRID: SCR_021135. Purchase of the Zeiss 880 confocal microscope used in this research was made possible through the National Institutes of Health s10 Grant 1S10OD23528-01. Portions of the text in this manuscript were edited solely for clarity and grammar using ChatGPT (GPT-4 Turbo, OpenAI); no content or scientific interpretation was generated by the language model.

## Ethics Statement

The animal study was reviewed and approved by the University of California, Santa Cruz animal care and use committee (IACUC).

## Data availability statement

All requests for data, resources, materials, and additional information should be directed to the lead contact, Euiseok J. Kim. This study did not generate new biological reagents. Any additional data reported in this paper will be shared by the lead contact upon reasonable request.

